# SC-JNMF: Single-cell clustering integrating multiple quantification methods based on joint non-negative matrix factorization

**DOI:** 10.1101/2020.09.30.319921

**Authors:** Mikio Shiga, Shigeto Seno, Makoto Onizuka, Hideo Matsuda

## Abstract

Unsupervised cell clustering is important in discovering cell diversity and subpopulations. Single-cell clustering using gene expression profiles is known to show different results depending on the method of expression quantification; nevertheless, most single-cell clustering methods do not consider the method.

In this article, we propose a robust and highly accurate clustering method using joint non-negative matrix factorization (joint NMF) based on multiple gene expression profiles quantified using different methods. Matrix factorization is an excellent method for dimension reduction and feature extraction of data. In particular, NMF approximates the data matrix as the product of two matrices in which all factors are non-negative. Our joint NMF can extract common factors among multiple gene expression profiles by applying each NMF to them under the constraint that one of the factorized matrices is shared among the multiple NMFs. The joint NMF determines more robust and accurate cell clustering results by leveraging multiple quantification methods compared to the conventional clustering methods, which uses only a single quantification method. In conclusion, our study showed that our clustering method using multiple gene expression profiles is more accurate than other popular methods.

## 1 Introduction

Single-cell gene expression analysis measures gene expression on a cell-by-cell basis and analyses the functions and properties of the cell using this information. The gene expression levels measured for each cell are recorded as a matrix called the “gene expression profile”. Unsupervised cell clustering using gene expression profiles is one of the fundamental single-cell gene expression analyses. It is a useful method for the classification and identification of unknown cell groups and for discovering the diversity and subpopulation of known cell types.

One of the objectives of single-cell analysis is to identify cell types by applying clustering and to extract a group of characteristic genes for a specific cell type as marker genes. However, single-cell analysis involves some differences (e.g., different gene expression values derived from different phases in the cell cycle and errors in measurement such as missing values) among cells, even if those cells were the same cell type. Therefore, conventional cell clustering is unsuitable for single-cell analysis. Additionally, the gene expression profile matrix often has a large number of rows and columns (cells and genes); thus, cell clustering requires feature selection or dimension reduction of the data. For these reasons, we need a robust and data-driven clustering method for features that reflect individual cell biases.

SC3 ([1]) and Seurat ([2]) are useful data-driven analysis tools for single cells. SC3 is a clustering method using an ensemble of multiple analysis results by PCA and k-means. Seurat is a method to analyze single cells, and Louvain clustering is used in the cell clustering. Matrix factorization, as a method of unsupervised learning, is another efficient method for cell clustering and is excellent in data dimension reduction or the extraction of latent features. In particular, non-negative matrix factorization (NMF) ([3]) is a suitable method for the dimension reduction to extract the features of gene expression profiles because NMF interprets the data as a superposition of the gene functions and cell characteristics. NMF factorizes a matrix into multiple matrices (basis and coefficients) under the constraint that all elements are non-negative, the product of those matrices constructs an approximate matrix of the input matrix. NMF is applied to various real data because of its non-negative feature and has been adapted to microarray data for clustering or feature extraction ([4]; [5]; [6]; [7]; [8]). We can analyse more details of genes and cells (e.g., finding marker genes and performing unsupervised cell clustering) with the feature matrices extracted by performing NMF.

Single-cell analysis is a series of procedures that consists of a number of methods. Multivariate analysis such as clustering needs quantification of gene expression, which converts the output of the sequencer into gene expression counts. Quantification is a critical factor for the subsequent analysis. Different quantification methods make the different quantified gene expression counts even when the same RNA-Seq reads are used. The greatest difference in quantification methods is whether the method is based on the genome references or transcriptome indices. For example, in methods based on the references, RNA-Seq reads are mapped to a reference genome using genome mapping tools such as Bowtie2 ([9]) and STAR([10]). Then, the gene expression counts are obtained from the results using expression estimation tools such as RSEM([11]) and Cufflinks([12]). In contrast, alignment-free quantification tools such as Salmon and kallisto quantify gene expression counts from RNA-Seq reads without mapping to a genome reference. To date, many methods for quantifying gene expression from RNA-Seq reads have been proposed; however, a consensus has not yet been reached on the best quantification method for all data ([13]; [14]). Moreover, since each quantification method has different measurable genes, the conventional analysis methods for cell clustering using gene expression quantified by only one method are strongly biased.

We usually observe that different quantification pipelines produce similar but different gene expression profiles even when we apply them to the same RNA-Seq reads. Since the cells included in the data are exactly the same, it is desirable to decompose the features with biases depend on each quantification method and the common factors for cell clustering. Our technical challenge is that how we can integrate the different quantification method. In this article, we propose SC-JNMF, a novel unsupervised clustering method using NMF to consider the differences of quantification methods and extract the common factors over multiple gene expression profiles. Our method achieved highly accurate cell clustering compared to other popular methods and showed usefulness for finding marker genes.

## 2 Methods

### 2.1 Outline of SC-JNMF

Gene expression profiles generated with different methods have different genes and expression values due to the biases of the methods, even if the profiles are generated from the same RNA-Seq reads. However, the information about genes and cells common to these profiles should be the same because of the shared origin. Furthermore, the results of cell clustering using these gene expression profiles should be the same. That is, in gene expression profiles generated from the same RNA-Seq reads by different methods, different features derived from the difference in methods and shared features derived from the original RNA-Seq reads should be extracted.

We proposed SC-JNMF, a method that extracts latent features from different gene expression profiles at the same time by using joint NMF, which can express matrix data as the product of lower-dimensional matrices, one of which is shared. SC-JNMF extracts the latent features in different gene expression profiles by a similar approach to NMF and uses them for cell clustering and gene analysis.

We show the outline of our method in Fig 1. We created two different gene expression profiles by different quantification methods from the same RNA-Seq reads. Then, we extracted the common factor using jointed NMF and extended it to perform multiple NMFs in parallel with two different basis factors ***W***_1_, ***W***_2_(derived from the different methods) and shared factors ***H*** (derived from the original RNA-Seq reads). Finally, we perform cell clustering by using these extracted features and any appropriate clustering methods, such as hierarchical clustering.

**Figure 1:**
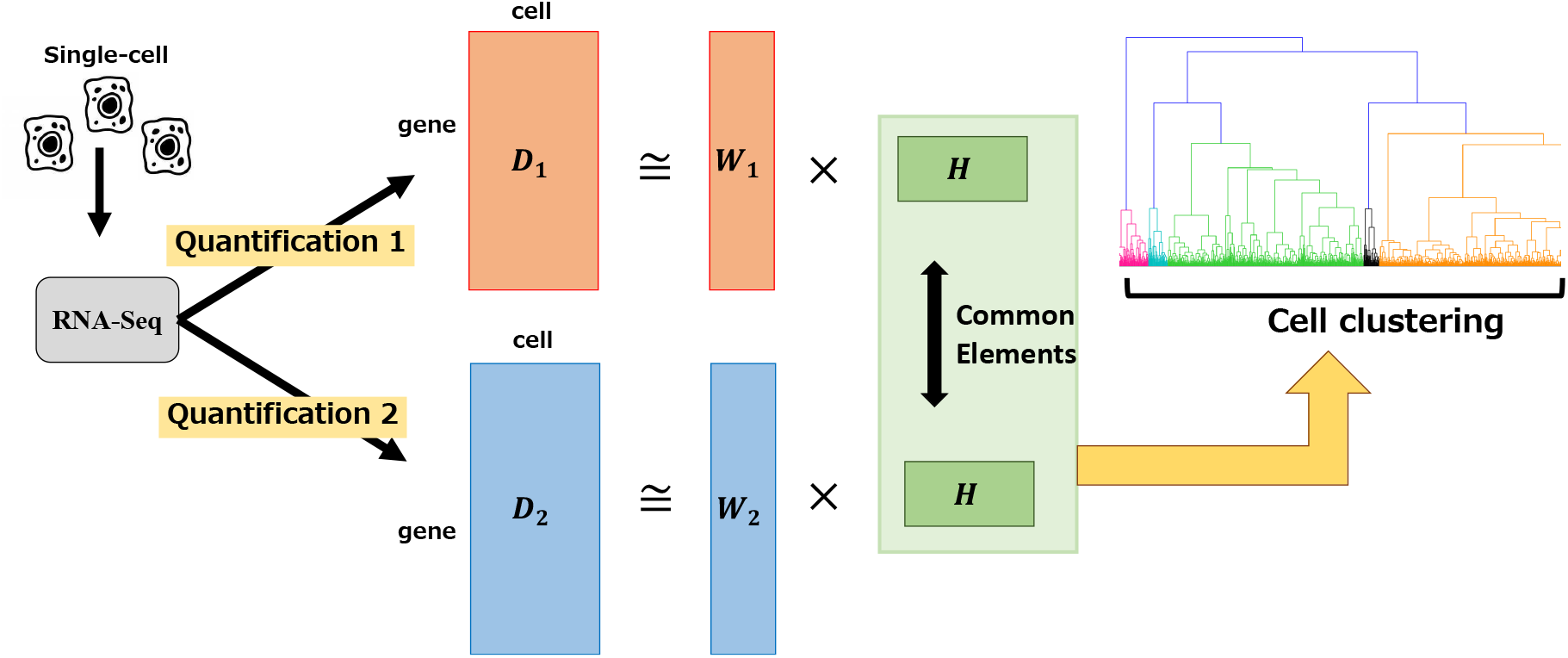
A workflow of SC-JNMF. (i) Make multiple gene expression profiles using different quantification method. (ii) Extract the common factor among these gene expression profiles from the same RNA-Seq reads. (iii) Cell clustering using hierarchical clustering with extracted common factor.

### 2.2 Joint non-negative matrix factorization

We consider the given data as a non-negative matrix 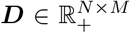. Non-negative matrix factorization finds the basis matrix 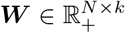 and coefficient matrix 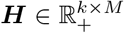, where all the elements are non-negative, so that these matrix product approximate the input data matrix.

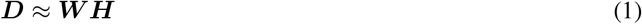

NMF minimizes the distance between matrix ***D*** and matrix ***WH***. Here, we consider the distance as the Euclidean norm. Thus, the objective function that NMF minimizes is as follows:

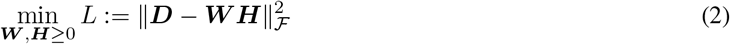

Here, we consider a simultaneous matrix factorization of two matrices using NMF. Given the two input data as non-negative matrices 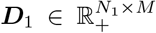 and 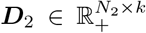, joint NMF finds the basis matrices 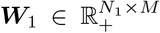, 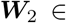 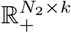 corresponding to each approximated input matrix and the common coefficient matrix 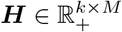, minimizing distances between the given matrices and the approximated matrices. In our proposed method, we add the L1 norm constraint to this objective function so that the factorized matrix is sparse. Thus, SC-JNMF finds the matrix ***W***_1_, ***W***_2_, ***H*** that minimizes the following objective function

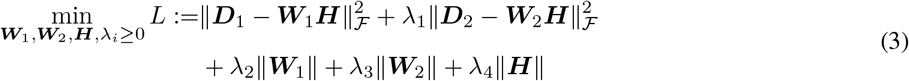

where, λ_*i*_(*i* = 1, …, 4) are regularization parameters.

By applying Jensen’s inequalities to the first and second terms of the objective function, the function to be minimized can be rewritten as follows.

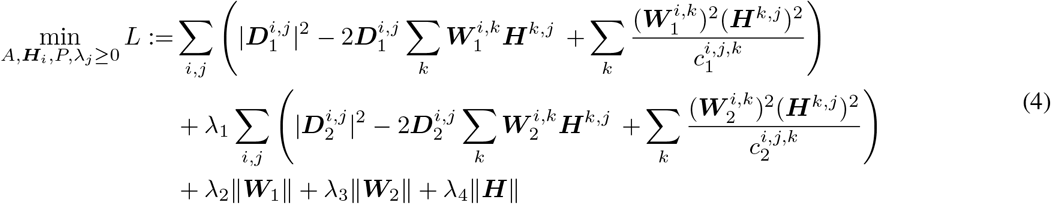

where,

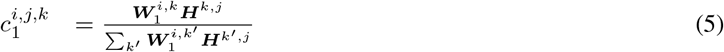

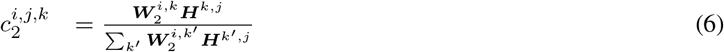

We find the each element of ***W***_1_, ***W***_2_ and ***H*** that minimizes the objective function by performing partial differentiation for them. Thus, the variable updates become

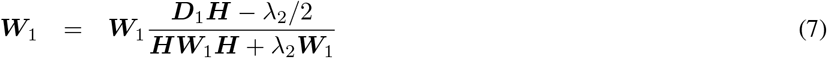

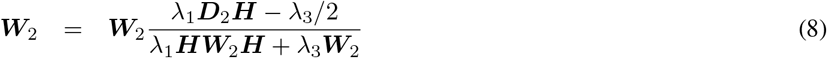

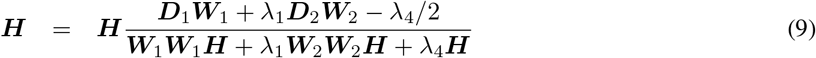

### 2.3 Applications

#### 2.3.1 Cell clustering

The extracted matrices can be used to perform highly accurate clustering. In our proposed method, we used hierarchical clustering (Ward’s method), which is one of the typical unsupervised clustering methods used for cell clustering. In this study, the adjusted Rand index (ARI) is used to evaluate the performance of clustering.

Given an *n* object set *S* = {*O*_1_, …, *O*_*n*_}, and suppose *U* = {*u*_1_, …, *u*_*R*_} and *V* = {*υ*_1_, …, *υ*_*C*_} represent two different partitions of S. The Rand index is defined by the following formula:

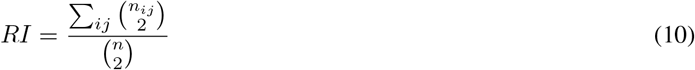

where, 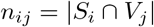, 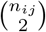 is binomial coefficient calculated as *n*_*ij*_(*n*_*ij*_ − 1)/2.

The Rand index calculates the ratio of the total of

i. the number of pairs are placed in the same class in U and the same class in V
ii. the number of pairs are placed in the different classes in U and in different classes in V

to the total number of pairs in set S.

In addition, the ARI considers the results of random clustering. The ARI is defined by the following formula:

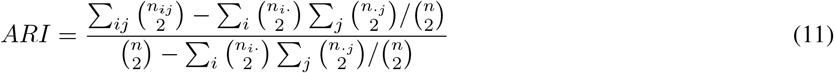

where, *n*_*i*·_ = Σ_*j*_*n*_*ij*_, *n*_·*j*_ = Σ_*i*_*n*_*ij*_.

#### 2.3.2 Parameters estimation

NMF is evaluated based on the value of the objective function to be minimized (the difference between the original matrix and the reconstructed matrix). However, the optimization problem of NMF is NP-hard, and only its convergence in minimizing the objective function is guaranteed ([15]). Therefore, there are several different local minima that can be obtained by NMF. To mitigate these problems, the factor’s sparseness in NMF is an important property in that it is a sufficient condition for the uniqueness of the solution, and we consider the sparseness of the factor in the evaluation of the proposed method and in the choice of parameters. In this study, the sparseness of the matrix 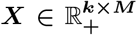 is formulated as proposed by Hoyer([16]).

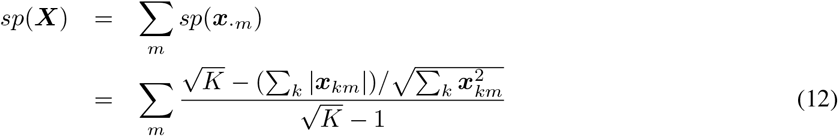

where, *K* is the rank of ***X***.

#### 2.3.3 Gene analysis

Our proposed method can perform not only cell clustering but also gene analysis by using the extracted features. The factors in the coefficient matrix that show higher values in specific clusters than in other clusters indicate latent features of the cluster. Therefore, the same factors in the basis matrices also show latent features of the cluster, and the genes that show higher values in the basis matrices reflect characteristic features of the cluster, in other words, marker genes.

## 3 Results

To assess the accuracy of our method, we performed cell clustering with 4 datasets. In this clustering, we estimated the optimal parameters (the number of ranks in matrix factorization) using the trade-off relationship between sparseness and loss. Additionally, we analysed more details of the factorized matrices by showing the relevance to marker genes.

### 3.1 Dataset

We used 4 different mouse/human single-cell datasets (Table 1). including RNA-Seq reads and the quantified gene expression levels measured by each method in previous studies. In addition, we newly quantified the gene expression levels in each cell from the RNA-Seq reads using Salmon(v1.0.0) and GENCODE (Mouse Release 21/ Human Release 32). In advance, we removed any cell types that were unclear in previous studies.

**Table 1:** Dataset and quantification method in previous studies.

| Dataset | Treutlein | Pollen | Segerstolpe | Xin |
| --- | --- | --- | --- | --- |
| Tissue/process | Lung epithelium | Brain | Pancreas | Pancreas |
| Quantification | TopHat v2.0.8, Cufflinks v2.0.2 | RSEM v1.2.4, TopHat v2.0.4 | STAR v2.3.0e, rpkmforgenes | CLC Bio Genomics Workbench v7.0 |
| Reference | mm10 | hg19 | hg19 | GRCh37 |
| The number of cells | 80 | 259 | 2166 | 1492 |
| The number of classes | 5 | 10 | 12 | 4 |
| Citation | [18] | [19] | [20] | [21] |

As preprocessing before performing joint matrix factorization, we applied a gene filter to the input data, as in the SC3 method ([1]), and then log2 transformed the data. Finally, we normalized the data so that the sum of each gene *L*^1^ norm was 1.

### 3.2 Rank estimation using loss and sparseness

There has been much discussion on the estimation of rank number in NMF; however, no consensus has been reached yet. In addition, our method has another hyperparameter, the regularization parameter on the term of L1 norm. Regarding the optimization of NMF, the lower loss and the higher sparseness(L1/L2) is better. Fig 2 shows the transition of ARI, sparseness, and loss with different ranks in joint NMF for all datasets. In general, the loss decreases and the sparseness increases as the rank increases. Therefore, the rank number that stops the decrease in loss or increase in sparseness is optimal because the factorized matrices with larger rank number include the information that is not necessary to represent the principal features of the input matrix. In this experiment, the Treutlein and Pollen datasets show the rank number that stops the increase in sparseness (4-6 in the Treutlein dataset, 7-9 in the Pollen dataset, 15-30 in the Xin dataset); however, the Segerstolpe dataset showed only a monotonic transition of sparseness and loss. We also ran the with different λ_4_ (the regularization parameter for the term on sparseness of matrix *H*). The differences in sparseness were confirmed in the Treutlein and Pollen datasets, while the Segerstolpe and Xin datasets showed almost the same sparseness in each rank.

**Figure 2:**
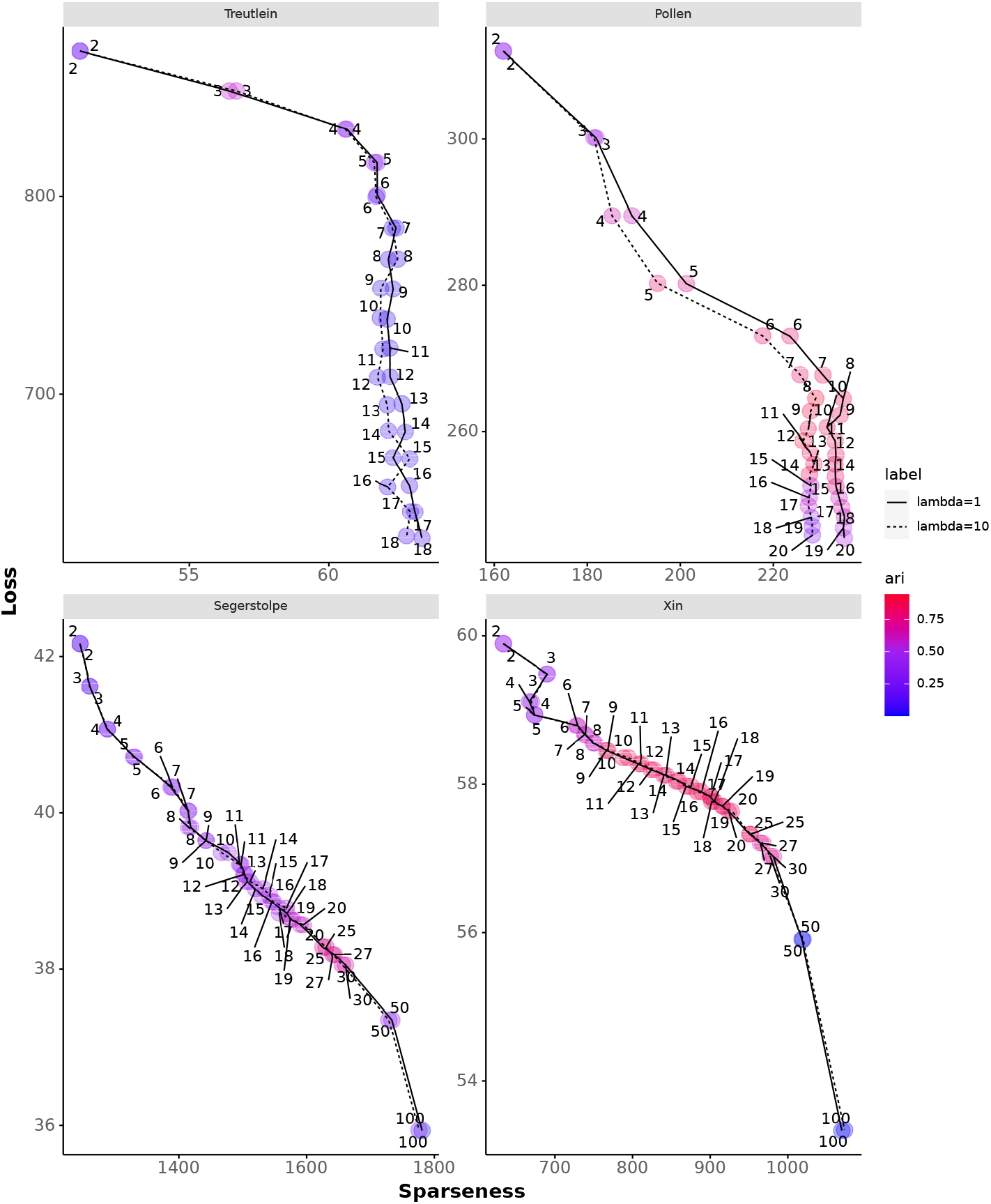
Loss and Sparseness of matrix ***H*** in each ranks. we ran the experiment in two way with different regularization parameters (λ_4_ = 1 and λ_4_ = 10).

### 3.3 The accuracy of cell clustering

We compared the accuracy of cell clustering using our proposed method to that obtained using other major unsupervised clustering methods. LSNMF : One of the NMF methods using the projected gradient method by ([17]). SC3 : The clustering method using the results of an ensemble of multiple analyses by PCA and k-means proposed by ([1]). Seurat : The method to analyse single cells proposed by ([2]). For cell clustering, this tool mainly uses Louvain clustering.

Fig 3 shows the ARI score of the original (quantified in a previous study) and alternative (quantified using Salmon) data for each clustering method. In the LSNMF method, as in joint NMF, classification by hierarchical clustering was performed using a matrix of features of the cells. For LSNMF and SC3, each run was performed 10 times considering the efficiency of the initial value, and for Seurat, we plotted the highest ARI score in the runs as the resolution parameter was increased from 0 to 1 in steps of 0.1. For our proposed method, we performed 10 runs. Each run consists of 10 runs and extracts an ARI score with the top combined rank in terms of sparseness and loss (the top ranking of sparseness is minimum and the top ranking of loss is maximum). The rank number was determined as follows: Treutlein:*k* = 5, Pollen:*k* = 8, Segerstolpe:*k* = 25, Xin:*k* = 16, according to the results of our experiment described in the next section. In addition, we determined the regularization parameter λ_1_ = |*geneset*1|/|*geneset*2|, λ_2_ = 0, λ_3_ = 0. For these datasets, λ_2_ and λ_3_ did not have a good influence on the clustering; in other words, the regularization terms of |*W*_1_|, |*W*_2_| were excessively strong constraints. We ran a two-way experiment with λ_4_ = 1, λ_4_ = 10. Hierarchical clustering cannot determine the number of clusters; therefore, we set the same number reported in previous studies in advance.

**Figure 3:**
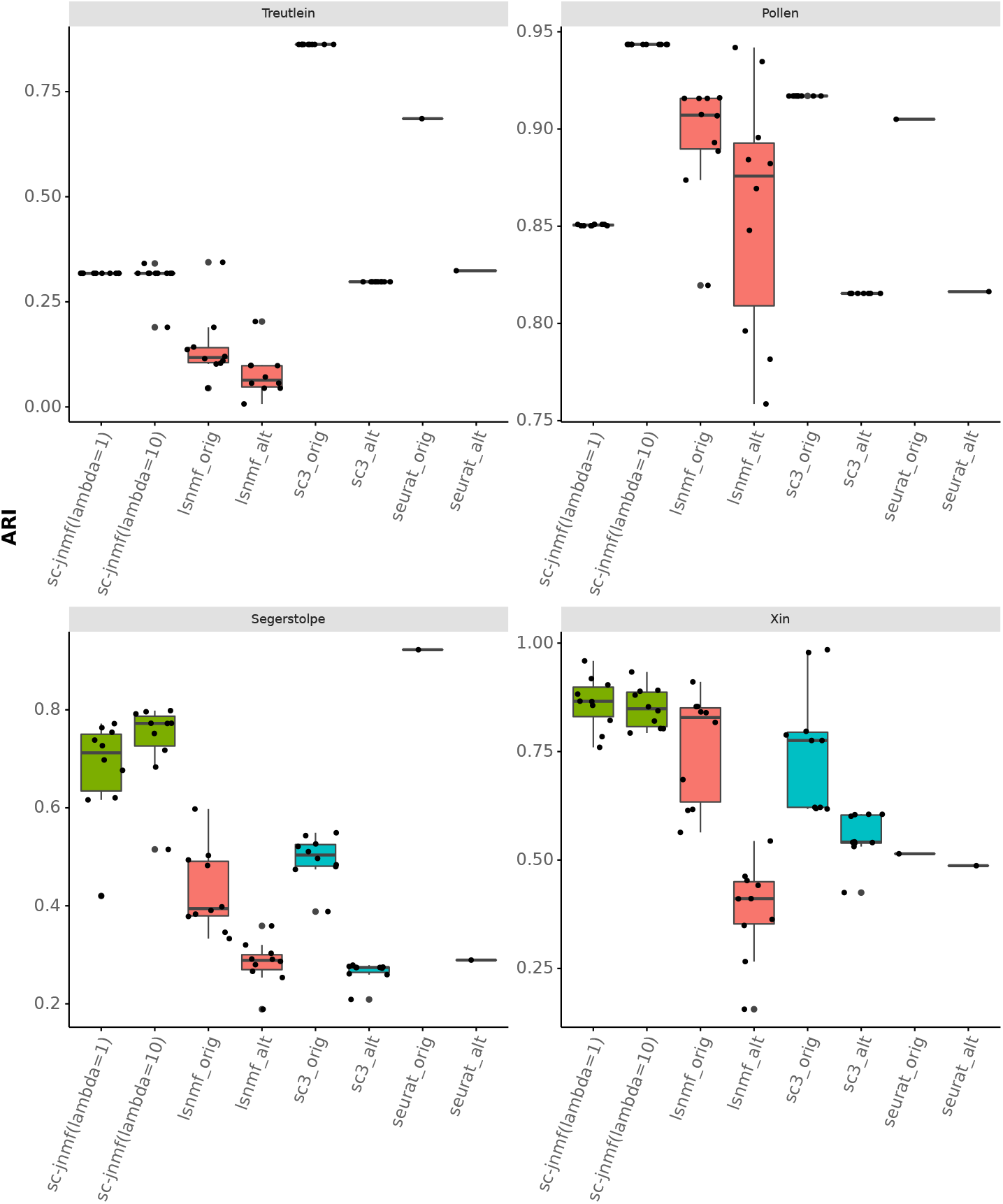
The accuracy (ARI) of cell clustering in each method. For comparison methods, we performed experiment with original gene expression profile (quantified in the previous research) and alternative gene expression profile (quantified using Salmon). For our proposed method, we ran the experiment in two way with different regularization parameters (λ_4_ = 1 and λ_4_ = 10).

In the Pollen and Xin datasets, our proposed method performed accurate cell clustering. On the other hand, in the Treutlein and Segerstolpe datasets, the ARI score of the proposed method was higher than that of LSNMF but not the highest score. Resulting clusters of our method is stable compared to conventional NMF method.

### 3.4 Gene analysis using factorized matrix

We found marker genes of the Xin dataset with the factorized matrices. The Xin dataset has data on 1492 single cells and 4 classes (alpha, beta, delta, PP) in the pancreas. We show the results in Fig 4. The factor of the coefficient matrix shows some characteristic patterns in each cell cluster Fig 4A. Some factors show marked features for the cell groups (e.g. factor 2 and factor 5 show higher values in delta and PP cells). Next, we calculated the correlation of factors between the basis matrices in the common genes (Fig 4B). The factors in the basis matrices showed similar features. We also show the values of marked factors for delta cells in the coefficient matrix in 4C. Almost all genes showed similar factor values between base1 and base2; however, we confirmed that some of the genes observed in either gene expression profile also had high values. We also show the marker genes of delta cells in this dataset (Fig 4D) in the scatter plot. The factor values of these genes tend higher than others regardless of whether the gene is observed in both gene expression profiles or only ether one.

**Figure 4:**
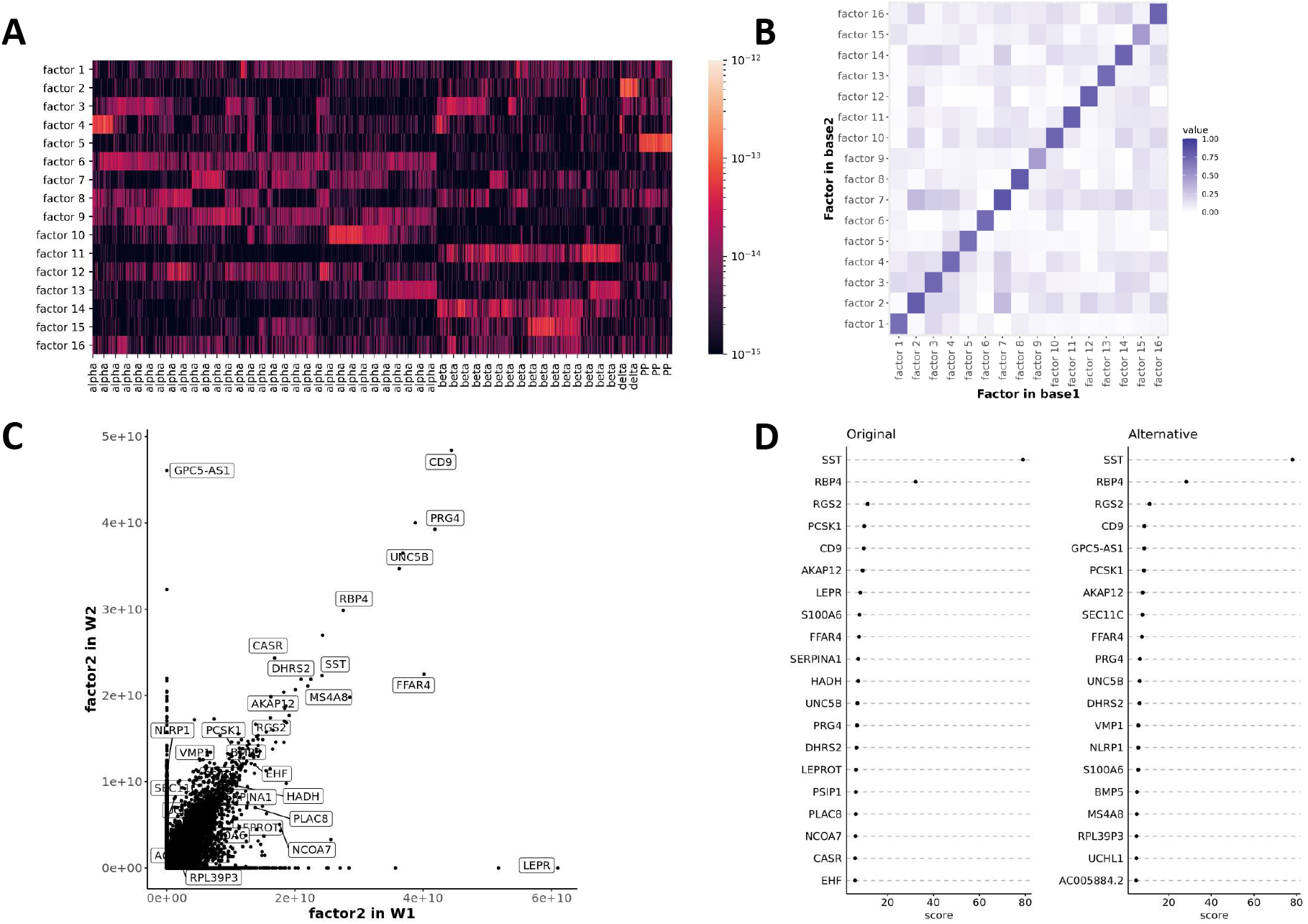
Gene analysis using our method in Xin dataset. (A) A heatmap of our coefficient matrix. (B) The correlation of the factor values in genes common to basis matrix1 and basis matrix2. (C) The “factor 2" values of each genes in base1 and base2. (D) Marker genes (differentially expressed genes) of delta cell in Original and Alternative gene expression profile detected by Scanpy. The score means the z-score underlying the computation of a p-value for each gene for delta cell.

## 4 Discussion

Conventional clustering methods have hyperparameters that we have to decide intuitively, and the NMF method also needs to determine the rank number. Previous studies analysed gene expression data using NMF and defined the rank number based on the number of cell clusters. However, this approach is not optimal for extracting the latent feature from given data. We defined the criteria for “good factorization” as the minimum loss and maximum sparseness. We provided a method to decide the hyperparameters of our method (the rank of factor matrices and the regularization parameter) considering increasing sparseness and decreasing loss. However, the optimal number of ranks for the Segerstolpe dataset is not obtained by deciding based on sparseness and loss. The optimal rank is not always clear depending on the dataset even with our provided method, so more reliable rank estimation that is applicable to all data sets is necessary.

Thus far, we showed the possibility that our unsupervised clustering method has high accuracy with using the differences in gene expression quantification methods. In comparison with previous studies, our proposed method showed highly accurate clustering results. However, the proposed method showed worse accuracy than SC3 in the original Treutlein datasets and Seurat in the original Segerstolpe datasets, and showed the differences in accuracy due to the differences in regularization parameters in the Pollen dataset. The Treutlein and Pollen datasets have fewer cells than the others, which is one of the most important features in our experiments. Dataset size is an important factor (common to machine learning approaches) in high accuracy, and this characteristic also applies to our method.

We presented more details about the factors of the factorized matrix and the relationships of marker genes. The coefficient matrix had characteristic factors in each cell cluster, and both basis matrices had similar factors between the matrices. The marker genes of a cell cluster have high values of the factor in the basis matrices that characterize a specific cell cluster in the coefficient matrix, regardless of observation in both gene expression profiles or only one; in other words, the factor values in the basis matrices that characterize a specific cell cluster in the coefficient matrix are related to the marker genes for the cluster. This result suggested that the genes showing high values in the basis matrices probably have some important features of the cluster with high values in the coefficient matrix. In particular, we should pay particular attention to those genes observed only in either gene expression profile because they are not considered in conventional methods.

## 5 Conclusion

We proposed SC-JNMF, which performs cell clustering using common factors extracted from multiple gene expression profiles quantified by different methods. As a result, it is possible to perform robust analysis compared with the case in which only a single quantification method is used because it is unnecessary to consider the differences in gene expression profiles obtained from different profiles. The accuracy (ARI) of cell clustering obtained using our method was higher than that of other major clustering methods. We also showed the transition of loss, sparseness and ARI in each rank, and the results suggested that the number of ranks in our method that stops the increase in sparseness tends to show the best ARI score. In addition, we showed the details of the extracted features with our method. The genes characteristic of specific cell groups (marker genes) showed remarkable values in the factors of factorized matrices; in other words, these results suggested a potential for identifying important genes in the dataset. Some genes may not be counted depending on the methods to use; they can be detected by our method if they are potential markers.

## References

[1] Vladimir Yu Kiselev, Kristina Kirschner, Michael T. Schaub, Tallulah Andrews, Andrew Yiu, Tamir Chandra, Kedar N. Natarajan, Wolf Reik, Mauricio Barahona, Anthony R. Green, and Martin Hemberg. SC3: Consensus clustering of single-cell RNA-seq data. Nature Methods, 14(5):483–486, 2017.

[2] Rahul Satija, Jeffrey A. Farrell, David Gennert, Alexander F. Schier, and Aviv Regev. Spatial reconstruction of single-cell gene expression data. Nature Biotechnology, 33(5):495–502, 2015.

[3] Daniel D Lee and H Sebastian Seung. Learning the parts of objects by non-negative matrix factorization. Nature, 401(October 1999):788–791, 1999.

[4] J.-P. Brunet, P. Tamayo, T. R. Golub, and J. P. Mesirov. Metagenes and molecular pattern discovery using matrix factorization. Proceedings of the National Academy of Sciences, 101(12):4164–4169, 2004.

[5] Chun Hou Zheng, To Yee Ng, Lei Zhang, Chi Keung Shiu, and Hong Qiang Wang. Tumor classification based on non-negative matrix factorization using gene expression data. IEEE Transactions on Nanobioscience, 10(2):86–93, 2011.

[6] Serena Nik-Zainal, Ludmil B. Alexandrov, David C. Wedge, Peter Van Loo, Christopher D. Greenman, Keiran Raine, David Jones, Jonathan Hinton, John Marshall, Lucy A. Stebbings, Andrew Menzies, Sancha Martin, Kenric Leung, Lina Chen, Catherine Leroy, Manasa Ramakrishna, Richard Rance, King Wai Lau, Laura J. Mudie, Ignacio Varela, David J. McBride, Graham R. Bignell, Susanna L. Cooke, Adam Shlien, John Gamble, Ian Whitmore, Mark Maddison, Patrick S. Tarpey, Helen R. Davies, Elli Papaemmanuil, Philip J. Stephens, Stuart McLaren, Adam P. Butler, Jon W. Teague, Goran Jönsson, Judy E. Garber, Daniel Silver, Penelope Miron, Aquila Fatima, Sandrine Boyault, Anita Langerod, Andrew Tutt, John W.M. Martens, Samuel A.J.R. Aparicio, Ake Borg, Anne Vincent Salomon, Gilles Thomas, Anne Lise Borresen-Dale, Andrea L. Richardson, Michael S. Neuberger, P. Andrew Futreal, Peter J. Campbell, and Michael R. Stratton. Mutational processes molding the genomes of 21 breast cancers. Cell, 149(5):979–993, 2012.

[7] Shihua Zhang, Chun Chi Liu, Wenyuan Li, Hui Shen, Peter W. Laird, and Xianghong Jasmine Zhou. Discovery of multi-dimensional modules by integrative analysis of cancer genomic data. Nucleic Acids Research, 40(19):9379–9391, 2012.

[8] Chunxuan Shao and Thomas Höfer. Robust classification of single-cell transcriptome data by nonnegative matrix factorization. Bioinformatics, 33(2):235–242, 1 2017.

[9] Ben Langmead and Steven L. Salzberg Fast gapped-read alignment with Bowtie 2. Nature Methods, 2012.

[10] Alexander Dobin, Carrie A. Davis, Felix Schlesinger, Jorg Drenkow, Chris Zaleski, Sonali Jha, Philippe Batut, Mark Chaisson, and Thomas R. Gingeras. STAR: Ultrafast universal RNA-seq aligner. Bioinformatics, 2013.

[11] Bo Li, Victor Ruotti, Ron M. Stewart, James A. Thomson, and Colin N. Dewey. RNA-Seq gene expression estimation with read mapping uncertainty. Bioinformatics, 2009.

[12] Cole Trapnell, Brian A. Williams, Geo Pertea, Ali Mortazavi, Gordon Kwan, Marijke J. Van Baren, Steven L. Salzberg, Barbara J. Wold, and Lior Pachter. Transcript assembly and quantification by RNA-Seq reveals unannotated transcripts and isoform switching during cell differentiation. Nature Biotechnology, 2010.

[13] Juliana Costa-Silva, Douglas Domingues, and Fabricio Martins Lopes. RNA-Seq differential expression analysis: An extended review and a software tool, 2017.

[14] Beate Vieth, Swati Parekh, Christoph Ziegenhain, Wolfgang Enard, and Ines Hellmann. A systematic evaluation of single cell RNA-seq analysis pipelines. Nature Communications, 2019.

[15] Stephen A Vavasis. On the complexity of nonnegative matrix factorization. Society for Industrial and Applied Mathematics Journal on Optimization, 20(3):1364–1377, 2009.

[16] Patrik O. Hoyer. Non-negative matrix factorization with sparseness constraints. Journal of Machine Learning Research, 5:1457–1469, 2004.

[17] Chin J. Lin. Projected gradient methods for nonnegative matrix factorization. Neural Computation, 19(10):2756–2779, 2007.

[18] Barbara Treutlein, Doug G. Brownfield, Angela R. Wu, Norma F. Neff, Gary L. Mantalas, F. Hernan Espinoza, Tushar J. Desai, Mark A. Krasnow, and Stephen R. Quake. Reconstructing lineage hierarchies of the distal lung epithelium using single-cell RNA-seq. Nature, 509(7500):371–375, 2014.

[19] Alex A. Pollen, Tomasz J. Nowakowski, Joe Shuga, Xiaohui Wang, Anne A. Leyrat, Jan H. Lui, Nianzhen Li, Lukasz Szpankowski, Brian Fowler, Peilin Chen, Naveen Ramalingam, Gang Sun, Myo Thu, Michael Norris, Ronald Lebofsky, Dominique Toppani, Darnell W. Kemp, Michael Wong, Barry Clerkson, Brittnee N. Jones, Shiquan Wu, Lawrence Knutsson, Beatriz Alvarado, Jing Wang, Lesley S. Weaver, Andrew P. May, Robert C. Jones, Marc A. Unger, Arnold R. Kriegstein, and Jay A.A. West. Low-coverage single-cell mRNA sequencing reveals cellular heterogeneity and activated signaling pathways in developing cerebral cortex. Nature Biotechnology, 32(10):1053–1058, 2014.

[20] Asa Segerstolpe, Athanasia Palasantza, Pernilla Eliasson, Eva Marie Andersson, Anne Christine Andréasson, Xiaoyan Sun, Simone Picelli, Alan Sabirsh, Maryam Clausen, Magnus K. Bjursell, David M. Smith, Maria Kasper, Carina Ämmälä, and Rickard Sandberg. Single-Cell Transcriptome Profiling of Human Pancreatic Islets in Health and Type 2 Diabetes. Cell Metabolism, 2016.

[21] Yurong Xin, Jinrang Kim, Haruka Okamoto, Min Ni, Yi Wei, Christina Adler, Andrew J. Murphy, George D. Yancopoulos, Calvin Lin, and Jesper Gromada. RNA Sequencing of Single Human Islet Cells Reveals Type 2 Diabetes Genes. Cell Metabolism, 24(4):608–615, 10 2016.

